# Growth phase-dependent responses of an *in vitro* gut microbiome to metformin

**DOI:** 10.1101/2020.02.05.936500

**Authors:** Zikai Hao, Leyuan Li, Zhibin Ning, Xu Zhang, Janice Mayne, Kai Cheng, Krystal Walker, Hong Liu, Daniel Figeys

## Abstract

*In vitro* gut microbiota models are often used to study drug-microbiome interaction. Similar to culturing individual microbial strains, the biomass accumulation of *in vitro* gut microbiota follows a logistic growth curve. Current studies on *in vitro* gut microbiome responses introduce drug stimulation during different growth stages, e.g. lag phase or stationary phase. However, *in vitro* gut microbiota in different growth phases may respond differently to same stimuli. Therefore, in this study, we used a 96-deep well plate-based culturing model (MiPro) to culture the human gut microbiota. Metformin, as the stimulus, was added at the lag, log and stationary phases of growth. Microbiome samples were collected at different time points for optical density and metaproteomic functional analysis. Results show that *in vitro* gut microbiota responded differently to metformin added during different growth phases, in terms of the growth curve, alterations of taxonomic and functional compositions. The addition of drugs at log phase leads to the greatest decline of bacterial growth. Metaproteomic analysis suggested that the strength of the metformin effect on the gut microbiome functional profile was ranked as lag phase > log phase > stationary phase. Our results showed that metformin added at lag phase resulted in a significantly reduced abundance of the Clostridiales order as well as an increased abundance of the *Bacteroides* genus, which was different from stimulation during the rest of the growth phase. Metformin also resulted in alterations of several pathways, including energy production and conversion, lipid transport and metabolism, translation, ribosomal structure and biogenesis. Our results indicate that the timing for drug stimulation should be considered when studying drug-microbiome interactions *in vitro*.

## Introduction

A major recent paradigm shift in medicine is the recognition of the important roles that the gut microbiome plays in drug therapy^1, 2^. The activity of drugs is usually dictated not only by the host, but often also by the gut microbiome. Evidences suggest that drugs can affect the functions and composition of the gut microbiome^3–5^. Increasing studies are focused on drug-microbiome interactions^1, 6^. *In vitro* gut microbiome culturing models are rapid, and cost-efficient approaches for studying drug-microbiome interactions and are increasingly applied to microbiome studies^7–11^. For example, the SHIME reactor (Simulator of the Human Intestinal Microbial Ecosystem)^11,12^, and the M-SHIME^13^ have been used to study drug effects on microbiome; Jalili-Firoozinezhad et al.^7,14^ have developed a microfluidic intestine-on-a-chip to study microbiome-related therapeutics, probiotics and nutraceuticals; Shah et al.^8^ have also presented a modular, microfluidics-based model (HuMiX, human-microbial crosstalk), which allows co-culture of human and microbial cells under conditions representative of the gastrointestinal human-microbe interface. Wu et al.^10^ have used an *in vitro* simulated human intestinal redox model (SHRIM) to investigate metformin-microbiota interactions directly. We have previously developed a 96-deep well plate-based culturing model (MiPro) facing high-throughput drug screening^15^.

Previous studies have added drug stimulation to the *in vitro* gut bacteria at various points during the growth phases of the gut microbiome. For example, Maier et al.^16^, Li et al.^15^ and Wang et al.^17^ studied microbial responses to drugs by inoculating bacterial strains or fecal inoculum into culture mediums containing drugs. Wu et al.^10^ used the SHRIM to explore the effect of metformin on a stabilized gut microbial community in vitro, and added the metformin after one week of stabilization in SHRIM. Similarly, with the SHIME models, compound are also added following 1~2 weeks of stabilization^11, 12^. Notably, bacteria can respond differently to environmental stimulations during different points in the growth phases. Early studies on the mechanism of action of penicillin showed that the bactericidal activity of β-lactam antibiotics depends on bacterial growth^18, 19^. This is best illustrated by the inability of penicillin to kill nongrowing bacteria in contrast to the rapid killing and lysis of the same cells during exponential growth^20^. We therefore postulated that the *in vitro* gut microbiome may also functionally respond differently to drugs added at different points in the growth phases, which, to the best of our knowledge, has not been reported yet.

In this study, we cultured a human gut microbiome using the Mipro model and added metformin at the lag, log and stationary phases of growth. Cultured samples were collected overtime for biomass and metaproteomic analyses. By analyzing growth curves and taxon-specific functional profiles, we demonstrated that adding metformin at log phase led to the greatest decline of bacterial growth while at the lag phage it induced the greatest changes on the functional profile. This highlights the importance of drug stimulation time in the investigation of drug-microbiome interactions.

## Results

### Determination of the microbiome growth curve

In this study, we hypothesized that drugs added at different growth phase of an in vitro microbiota may have different effects (Figure 1A). In order to determine the time points (i.e. lag, log and stationary phases) of drug stimulation, we first measured the growth curve of the *in vitro* gut microbiome. Briefly, a human gut microbiome was cultured in each well of a 96-well plate using MiPro^15^. Samples were taken at 15 time points in triplicates from 0-48 hours of culturing (Figure 1B). Optical density of each sample was measured at 595 nm (OD_595_), as a proxy of microbial biomass. The growth curve of the *in vitro* gut microbiome was calculated by fitting a logistic growth model to the data from each time point(Figure. 1B). The microbial biomass *Nt* at time *t* was described by the following logistic equation:

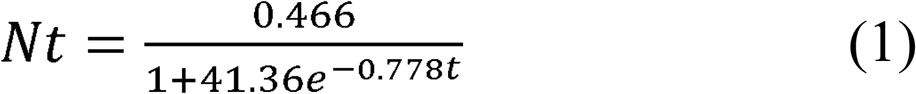

**Figure. 1.**
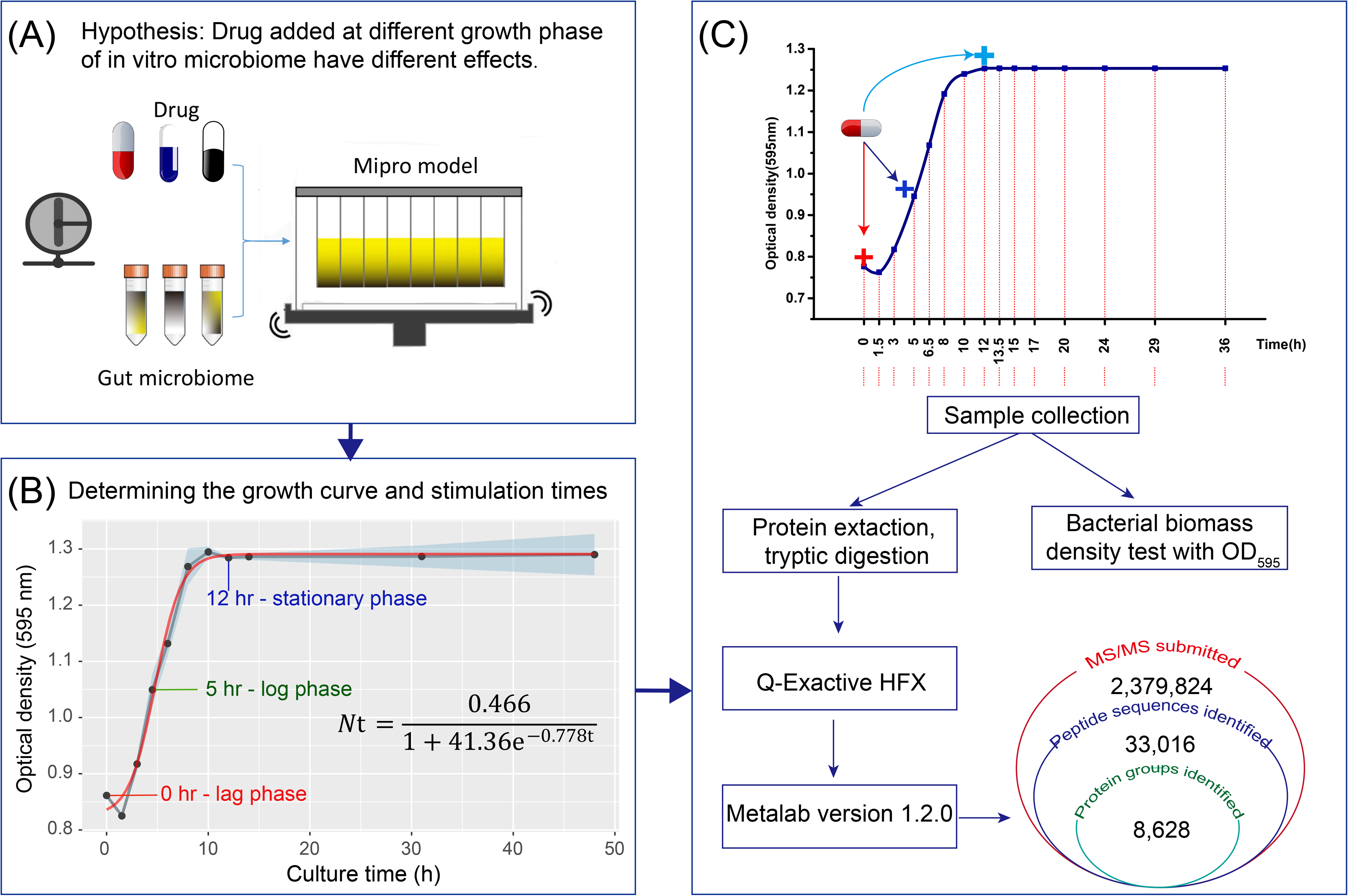
Experimental hypothesis and workflow. (A) Experimental hypothesis. (B) In vitro growth of the human gut microbiome in 48 hours. Black dots are means (n=3 individual wells) of the OD_595_ value of the culture, and blue ribbon stands for standard deviation. Red line is the logistic growth model fitted based on the mean OD_595_ value. (C) Experimental workflow. Human gut microbiome samples were cultured with metformin added at 0^th^ hour, 5^th^ hour and 12^th^ hour. Then, the samples in 96-well deep-well plates were collected for microbial biomass and metaproteomics analysis. Peptide and protein identification and quantification, taxonomic profiling, and functional annotation were performed using the automated MetaLab software^21^.

From Eq. (1) we calculated that the maximum possible microbial biomass *(K)* was 0.466, and the intrinsic growth rate of the microbiome (*r*) was 0.778. At 4.8 hr, the microbiome reached the maximum growth rate, i.e. the microbial biomass reached 1/2 *K*. The microbiome achieved stationary growth at about 12 hr.

We then tested the responses of the microbiome to metformin at three different times point representing different growth phases, i.e. 0 hr - lag phase (Lag), 5 hr - log phase (Log) and 12 hr - stationary phase (STA). Briefly, metformin was added at a final concentration of 7.6 mM into designated wells at the three growth phases^15^ (Figure 1C). Following each metformin treatment, samples were collected in triplicates over time for optical density and metaproteomic analyses (Figure 1C).

### Stimulation at the log phase resulted in the strongest antibacterial action

To compare the difference in growth curve between different treatment times, t-test and one-way ANOVA were performed. Results show that the biomass of the treatment groups (Lag/Log/STA) was significantly lower than the control group following the addition of metformin (Figure 2A). Moreover, after 20 hr, all metformin treatments showed loss of biomass while the control microbiome was still at the stationary phase, indicating that metformin has a significant antibacterial effect. As well, the trajectories of the curves were different between groups with the addition of metformin during the log phase causing a steep decline after 20 hours while the addition during the Lag and STA phases did not cause a significant continuous decline.

**Figure. 2.**
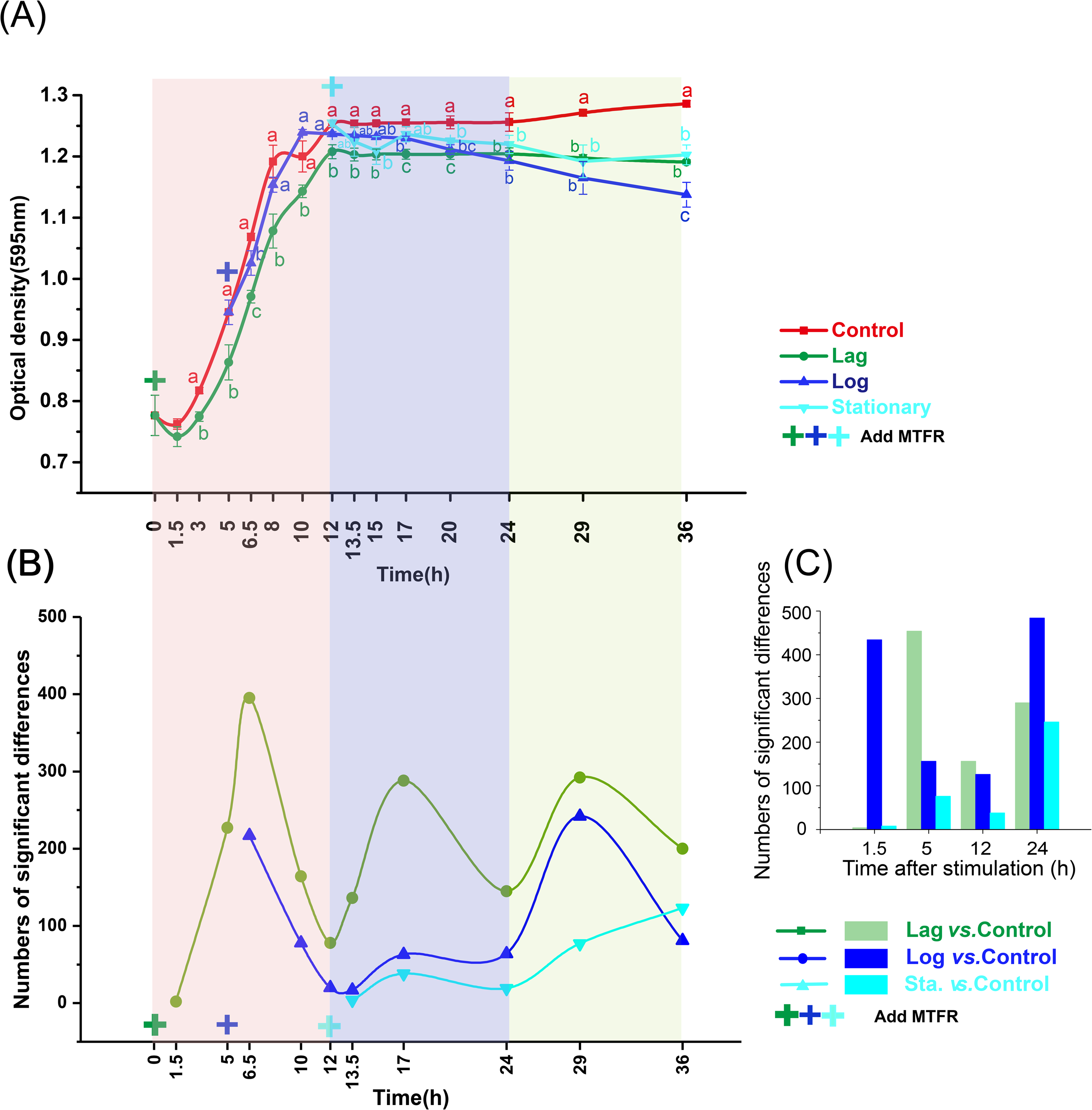
Metaproteomic response to metformin treatment was associated with the growth phase. To compare the difference in growth curve and metaproteomic profiles between different treatment times, t-test was performed at the time points for 1.5-5 hr (control vs. Lag only), and one-way ANOVA was performed at the time points for 6.5-36 hr. (A) Gut microbiome response differently to metformin on the growth curve when metformin added at different growth phase. Optical density at 595 nm (OD595) was used as a proxy of microbial biomass. Data were shown as averages and error bars represent SEM; statistically significant differences were shown with different letters (p < 0.05). T-test and one-way ANOVA with Duncan’s test was performed. (B) Numbers of significantly altered protein groups between the treatment group and control group changed over time (FDR-adjusted P<0.05). (C) Numbers of significantly altered protein groups between the treatment group and control group after the same hours of stimulation (FDR-adjusted P<0.05). T-test was performed.

### Microbiome functional response to metformin is associated with the microbial growth phase

We then explored whether adding metformin during the different phases can lead to difference in functional changes in the microbiome. Briefly, metaproteomic analysis was performed on 85 samples collected at different time points (5-7 time points in each group, and triplicates samples at each different time point, Supplementary Table S2). 2,379,824 MS/MS spectra were obtained from all 85 raw files, resulted in 33,016 peptides and 8,628 protein groups (FDR < 1%; Figure 1C). The average Pearson’s correlation coefficient between metaproteomic quality control samples was 0.992 ± 0.001 (mean ± SD, n= 3), and the correlation between each group of technical replicates ranged from 0.956 ± 0.028 to 0.990 ± 0.001 (n = 3; Supplementary Figure S1 and Table S1). These results indicated high reproducibility of both culturing and metaproteomics approach.

We first looked at the differences in protein expressions between treatment and control groups at each time point and after the same time of stimulation based on label-free quantification (LFQ) protein intensities (FDR-adjusted *P*<0.05). Different proteins had significant changes in expression at each time points and between treatment groups compared to the control group (Figure 2B) suggesting that the gut microbiome responded differently to metformin added at the three time points. Interestingly, introducing metformin at the lag phase led to a cycle of differentially expressed proteins with a first maximum at 6.5 hours and maximum about every 10-12 hours thereafter. During 24-36 hr, when metformin exerted significant antibacterial effects on the microbial biomass significantly declined (Figure 2A), the number of significantly altered proteins increased again. However, the Lag group had the highest response during the whole growth phase.

We then compared the number of significantly altered protein expression across the growth phases after an equal amount of time (1.5, 5, 12 24 hours) following the introduction of metformin (Figure 2C; FDR-adjusted *P*<0.05). The result confirmed that the introduction of metformin during different growth phases led to changes in protein expressions in the microbiome that remain distinct over time. Interestingly, the highest responses were observed while groups were in the log phase following treatment. After 24 hours of stimulation, the response in all treatment groups increased significantly, likely due to the significant antibacterial effect of metformin at that time.

The addition of metformin at different growth phase also led to alterations of microbiome genera over time (Supplementary Figure S3). Interestingly, the changes in the relative abundance of genera *Bacteroides* and *Fusobacterium* showed a cyclic pattern similar to the significantly altered proteins (Figure 2B), which may also explain the cyclic pattern observed in significantly altered proteins.

### Stimulation at the lag phase resulted in the strongest metaproteomic response

We further visualized the overall change of protein group intensities using principal component analysis (PCA) (Figure 3A). In most cases, the three groups were differentiated on the PCA score plots. Interestingly, the differentiation was related to the numbers of significantly altered protein groups at the corresponding points (Figure 2B). We further performed multivariate analysis of variance (MANOVA) to statistically compare metaproteomic differences between different groups. Hierarchical clustering was generated based on mahalanobis distances as a result of MANOVA (Figure 3C and Supplementary Figure S2). The MANOVA analysis showed that there were significant differences (p<0.05) between each treatment and the control at each time point (with exception of STA vs control at 13.5 and 24 hr). In particular, the PCA and MANOVA analyses suggested that the Lag group had the highest response among the three groups. PCA score plot (Figure 3B) and MANOVA analysis (Figure 3C) based on the last three time points (24, 29 and 36 hr) showed a clear separation between control, Lag, Log and STA groups, suggesting that the strengths of metaproteomic responses were ranked as Lag> Log > STA groups.

**Figure 3.**
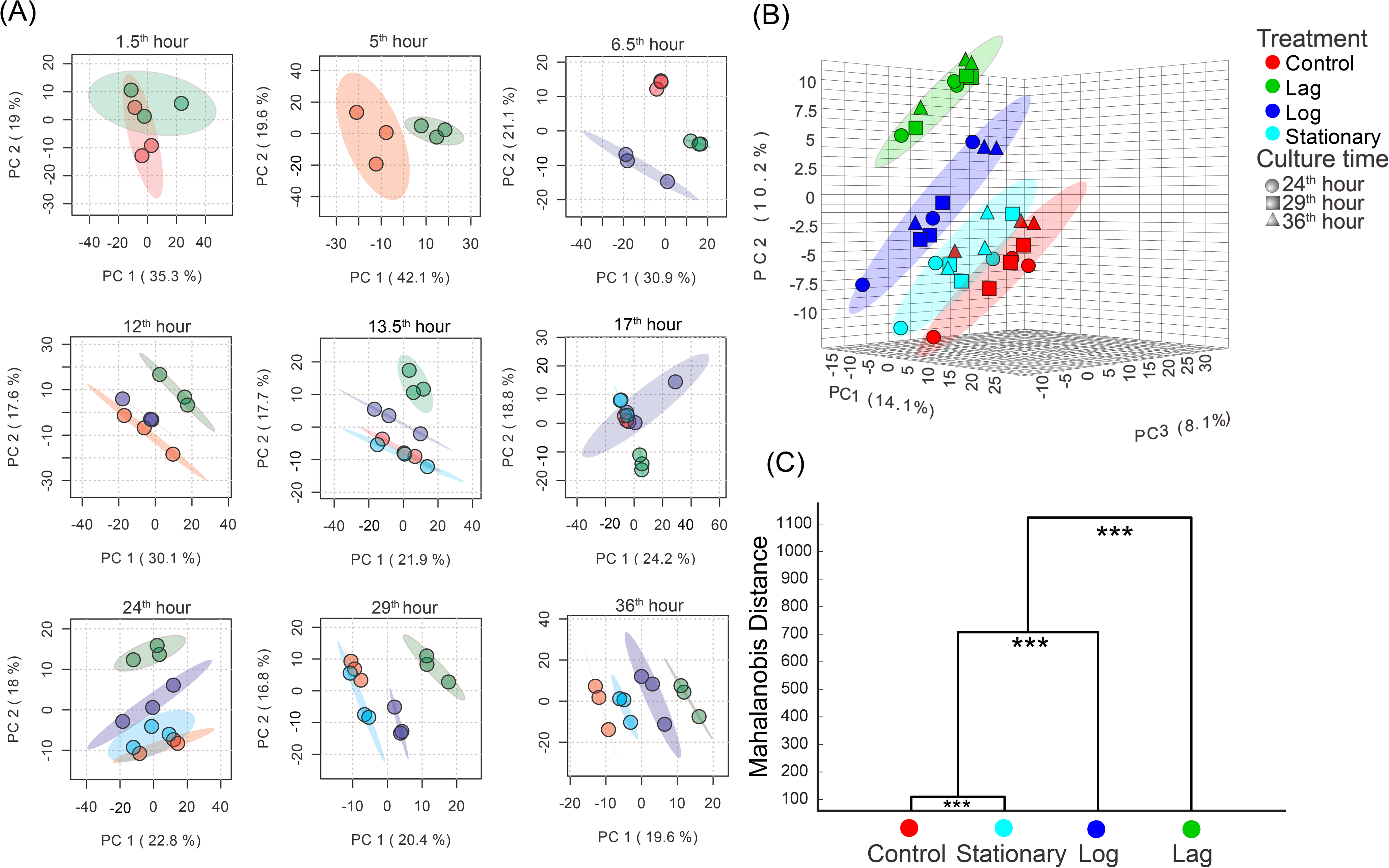
The alteration of gut microbiome in different groups response to metformin over time. (A) Principal component analyses (PCA) scores plots based on protein groups of the samples at the 1.5, 5, 6.5, 12, 13.5, 17, 24, 29 and 36 hr, respectively. (B) PCA plots based on the protein groups of the all samples during the 24-36 hr (24, 29 and 36 h). PCA plots showing metaproteomic profiles were different when metformin was added at different growth phases. (C) Clustering of different groups during the 24-36 hour based on mahalanobis distances calculated using MANOVA, ***, P < 0.001. The PCA plots and mahalanobis distances of MANOVA analysis showing strength of metformin effect on gut microbiome functional profiles was ranked as lag phase > log phase > stationary phase.

A time-series analysis using ANOVA - Simultaneous Component Analysis (ASCA) revealed patterns of protein expression that are time or treatment specifics as well as patterns that depend on both time and treatment simultaneously (interaction), Figure 4A-H. The pattern of “Treatment” or “Time” is the case where no time-treatment group interactions were modeled, only a treatment or time effect in all the groups. The score profiles of the first component (73.3% of variability) of the “Treatment” showed that the strength of the metformin effect on the gut microbiome was ranked as Lag group > Log group > STA group > Control group (Figure 2E). The score profiles of the first component (57.23% of variability) of the “Time” showed a negative effect through time for all groups (Figure 2F). The scores of all groups decreased rapidly during the log phase (1.5-12 hr) and gradually stabilized during the stationary phase (13.5-36 hr), which suggested that the strongest metaproteomic response to metformin treatment was associated with the growth phase (Figure 2F). The score profiles of the first two components of the “Interaction” (treatment plus time) described the different behavior for all groups over time (Fig. 2G-H). The score profiles of the first component (23.3% of variability) identified a marked effect through time for the Lag group which was different from the rest of groups. The Second component (10.35% of variability) identified a marked effect through time for the Log group which was different from the other groups. Finally, the scores of the STA group were similar to the control group in the score profiles of the “Treatment” and “Interaction”.

**Figure. 4.**
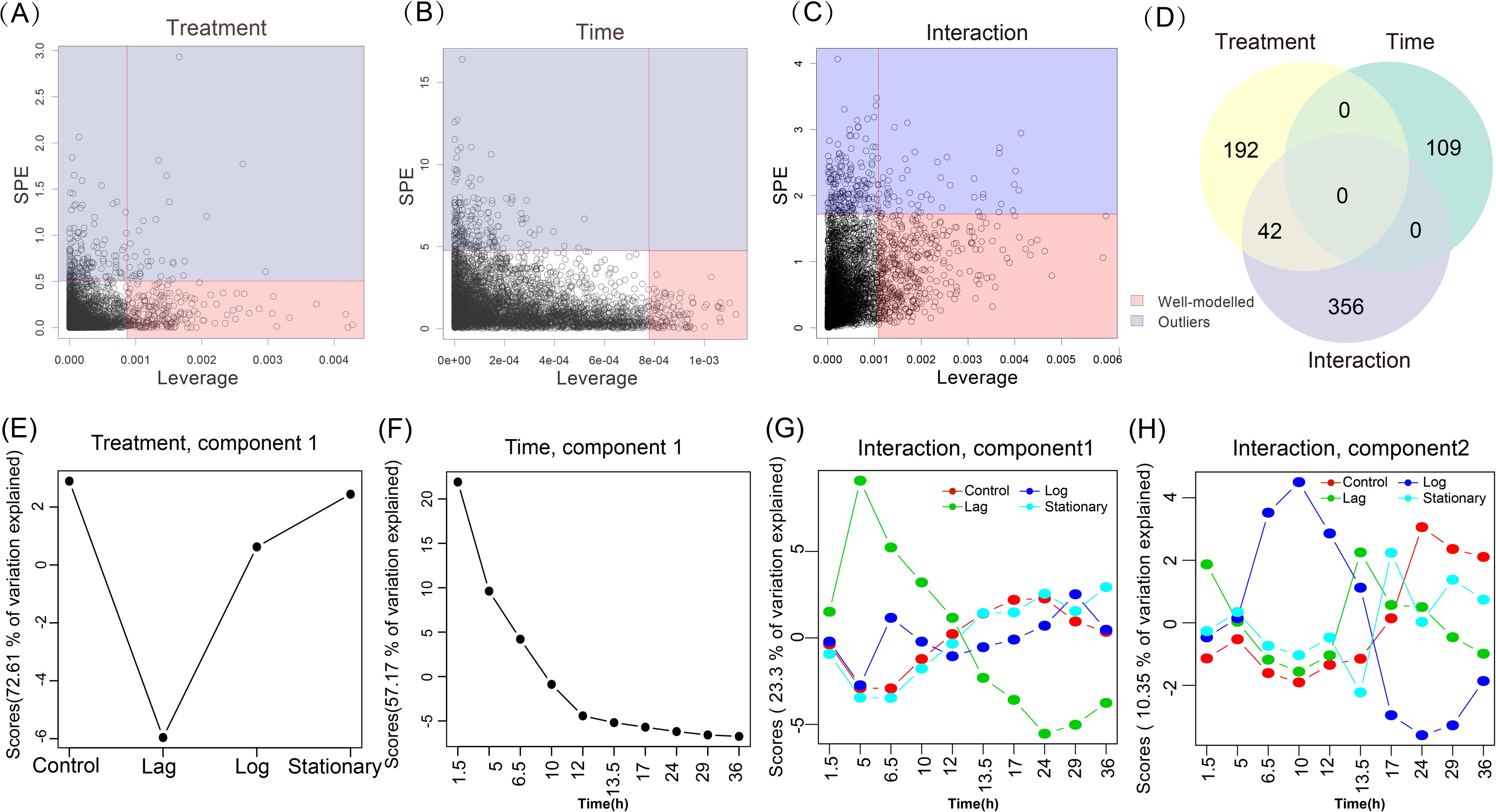
A time-series analysis using ANOVA - Simultaneous Component Analysis (ASCA). SPE and leverage statistics of the protein groups in “Treatment” (A), “Time” (B) and “Interaction” (C) by the ANOVA-Simultaneous Component Analysis (ASCA). Squared prediction error (SPE) and leverage cut-off values were indicated by horizontal and vertical lines, respectively. In ASCA analysis, most relevant (well modeled, pink filled area) protein groups in the ASCA-model will be those showing high leverage and low SPE. (D) Venn diagram showed, in total, 699 well modeled protein groups were identified, of which 234 in ‘Treatment’, 109 in ‘Time’ and 398 in ‘Interaction’, respectively. Additionally, 42 protein groups were identified in ‘Treatment’ and ‘Interaction’. Score profiles of component 1 of ‘Treatment’ (E) and ‘Time’ (F). Score profiles of the first two components of sub model ‘Interaction’ (Treatment versus Time) (G-H).

### Taxonomic source and functional profiles of altered proteins were distinct when stimulated at different growth phases

In ASCA analysis, the pattern of “Treatment” represents effect only due to metformin in all the groups. Therefore, to analyze the overall difference of taxon-specific functional responses of the gut microbiome stimulated at different growth phases, we selected well-modeled protein groups associated with ‘Treatment’ (with a high leverage and low SPE, Figure 4A) for detailed analysis. First, 234 well-modeled protein groups associated with ‘Treatment’ were identified (Supplementary Table S3). Then, hierarchical clustering and heatmap analysis revealed two different protein clusters (Figure 5A). Comparison of the expression profile of all protein groups in each cluster were shown in Figure 5B and E. In cluster I, the LFQ intensities of protein groups were lower in the Lag group as compared with the other treatment groups and the control (Figure 5B). In contrast, cluster II protein groups were higher in the Lag group (Figure 5E).

**Figure. 5.**
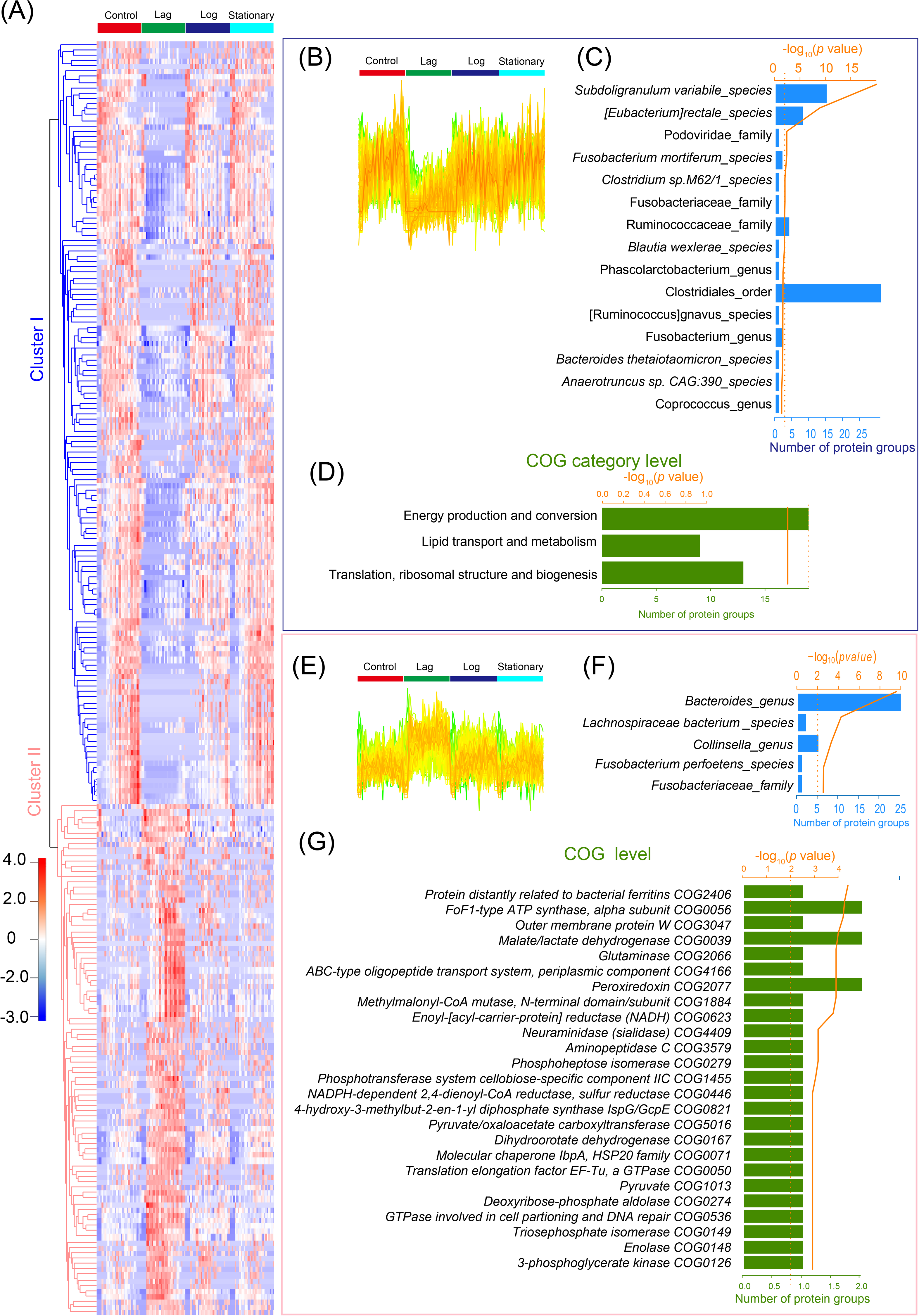
Metaproteomics revealed alterations of metaproteomic profiles when metformin added at different growth phases. (A) Heatmap of well-modeled protein groups associated with ‘Treatment’ by ASCA analysis. Protein groups with high leverage exhibit and low squared prediction error value are well-modeled. Heatmap shows the LFQ intensity of altered protein groups for each sample, 2 clusters (painted in blue and pink colors) were divided for further analyzed. (B, E) Alterations of protein groups in different groups in each cluster. (C, F) Alterations of taxonomic composition in each cluster. All the significantly taxonomy with a p-value cut off of 0.05 are shown. (D) Alterations of functional composition (COG categories) in cluster I. (G) Top 25 most significant altered COGs in cluster II. No significant COG category was enrichment in cluster II. The significantly altered COG categories and COGs with a p-value cut off of 0.05. Bar charts showed the total numbers of protein groups corresponding to each taxonomy or function (FDR-adjusted P<0.05).

We performed functional and taxonomic enrichment analysis of the protein groups of clusters I and II using the “Enrichment analysis” app from iMetaLab (https://shiny.imetalab.ca/). To identify the taxonomic source of these proteins, the representative protein for each protein group in the dataset were submitted to the app, and enriched taxa with a *p*-value cut off of 0.05 were obtained (Figure 5C and F). Significantly reduced proteins from the Lag group were enriched in order Clostridiales and family Fusobacterium, species *Subdoligranulum variabile, [Eubacterium]rectale, Fusobacterium mortiferum, Clostridium sp.M62/1* and *Blautia wexlerae;* while the increased proteins were enriched in genus *Bacteroides* and species *Lachnospiraceae bacterium* and *Fusobacterium perfoetens*. Interestingly, as shown in Figure S3, metformin added in the lag phase resulted in a sustained increase in the abundance of the genus *Bacteroides* after 10 hours of stimulation, which was different from the rest of treatment groups.

We then performed functional enrichment analysis based on the database of clusters of orthologous groups (COG). All the significantly altered COGs and COG categories with a *p*-value cut-off of 0.05 were shown in Figure 5D and Supplementary Table S6. At the COG category level, in the Lag group, energy production and conversion, lipid transport and metabolism, translation, ribosomal structure and biogenesis were significantly decreased, no COG category was significantly enriched by the increased proteins (Figure 5D and G). However, we identified 47 COGs from 14 COG categories that were significantly increased (Supplementary Table S6) in the Lag group. Functions related to the carbohydrate transport and metabolism (7 COGs), Translation, ribosomal structure and biogenesis (7 COGs) were among the most significantly increased COGs.

We further analyzed the taxon-specific functional responses of the gut microbiome in different treatment groups at 6.5, 17 and 36 hr (Supplementary Figures S4-S6). The results revealed that the gut microbiome responded differently to metformin added at different growth phases in each time point. For example, significantly altered protein groups at 36 hr were identified by one-way ANOVA with Fisher’s LSD test (FDR-adjusted p<0.05; Supplementary Table S4). Then, hierarchical clustering revealed five different protein clusters with >10 proteins (Figure S4A and B). Taxon and functional enrichment analysis were performed with the protein groups in each cluster (Figure S4C-D). To further analyze the taxon-specific functional difference in the three groups after the same number of hours of stimulation, we selected metaproteomic data after 12 and 24 hr of stimulation for taxon-specific functional analysis, as show in Supplementary Figure S7 and S8, respectively. The significantly upregulated and downregulated protein groups in treatment groups compared to controls were identified by *t*-test (FDR-adjusted p<0.05; Supplementary Table S5). Venn diagram indicates that after 12 or 24 hr of stimulation, the most of significantly altered protein groups in the different treatment group were unique. The metaproteomic results revealed that the gut microbiome responded differently to metformin added at different growth phases, even at the same time after stimulation. The uniquely altered proteins of different treatment group were from different taxonomy and function.

## Discussion

In this study, we evaluated whether adding drug treatment at different growth phase of *in vitro* human gut microbiome would have different impacts on the functions of the microbiome. Usually, *in vitro* bacterial growth involves four phases: lag phase, log (exponential) phase, stationary phase, and death (decline) phase^22, 23^. Bacteria adapt to the growth conditions in lag phase, which is the period the individual bacteria are maturing and not yet able to divide. The log phase is the period characterized by a logistic growth of bacterial biomass. The stationary phase results from a situation in which growth rate and death rate are equal, which is often due to a growth-limiting factor such as the depletion of an essential nutrient, and/or the formation of an inhibitory product such as an organic acid^24^. The death phase could be due to lack of nutrients and/or other injurious conditions^22, 23^.

In this study, we observed that the growth curve of the microbiome is similar to the growth of bacterial pure cultures^25^. We then performed three different treatments by adding metformin at the Lag, Log and stationary phases. We showed that the growth curves and metaproteomic profiles between treatment groups are significantly different. In agreement with previous reports^26, 27^, we also found that metformin has antibacterial action on *in vitro* gut microbiome (Figure 2A), especially after 24 hours of treatment. When subjected to nutrient limiting conditions, the bacteria become sensitive to metformin^28^. Stimulation at the log phase resulted in the strongest antibacterial action. It may well be that the antibacterial activity of metformin is dependent on the bacterial growth. The early studies on the mechanism of action of antibiotics showed that their bactericidal activity depends on bacterial growth^18, 19^, our study suggested similar response of the growing microbiota to metformin. Previous study indicated that metformin could significantly protect animals from tested pathogenic bacteria and it has potential as a noteworthy antibacterial agent to be used in human^27^. Our research indicated that bacterial growth should be considered in future studies of the antibacterial mechanism and activity of metformin.

We showed that the metaproteomic profiles varied over different stimulation time points, with the earlier stimulation leading to the greatest effects on the gut microbiome. During lag phase, bacteria adapt to the growth conditions, and synthesis of ATP, DNA, RNA, enzymes and other molecules occurs^22^. Adding metformin at lag phase may significantly affect the protein function in these biological processes. We showed that metformin added at lag phase significantly decreased functions of energy production and conversion, lipid transport and metabolism, translation, ribosomal structure and biogenesis, which is in agreement with several previous studies^5, 26, 28^. And metformin treatment also significantly increased the function about defense mechanisms, cell wall/membrane/envelope biogenesis. Most of these functions are closely related to the biological processes of bacteria in the lag phase^22^, which may explain why the metformin added at the lag phase resulted in the strongest metaproteomic response.

Several studies showed that metformin can alter gut microbiota composition^5, 26, 29, 30^, which also have been observed in our study. As with previous studies, our results indicated metformin causes a significantly increased abundance of the genus *Bacteroide*^29,31,32^, which was different from the rest of treatment groups. Metformin treatment at lag phase significantly increased defense mechanisms and cell wall/membrane/envelope biogenesis functions in the genus *Bacteroides*^33, 34^. The species of the genus *Bacteroides* have complex systems to sense and adapt to the growth environment, and multiple pump systems to expel toxic substances, they have the most antibiotic resistance mechanisms and the highest resistance rates of all anaerobic bacteria^33, 34^ Therefore, our study may explain why metformin added at lag growth phase can increase the abundance of the genus *Bacteroides*.

In summary, our results suggest that drugs can have different effects depending on the growth phase of the microbiome and that in vitro systems are useful to assess these effects. The effects of drugs on the microbiome included changes in microbial biomass, functional profiles and their taxonomic contributions. Our work revealed that metformin added during the earliest growth phase had the strongest functional/compositional effect on the microbiome. However, the addition of metformin at log phase led to the greatest decline of bacterial growth. It could also mean that the effects of metformin *in vivo* might be affected by the phase of the microbiome. This in vitro approach would also be useful to identify compounds that have effects in different growth phases providing alternative strategies to target the microbiome.

## Methods

### *In vitro* gut microbiome culturing and metformin treatments

Here, we used a 96-deep well plate-based culturing model (MiPro) reported in our previous studies^26^ for *in vitro* gut microbiome culturing. The medium composition for MiPro was based on our previously suggested medium composition^35^, which comprises: 2.0 g L^-1^ peptone water, 2.0 g L^-1^ yeast extract, 0.5 g L^-1^ L-cysteine hydrochloride, 2 mL L^-1^ Tween 80, 5 mg L^-1^ hemin, 10 μL L^-1^ vitamin K1, 1.0 g L^-1^ NaCl, 0.4 g L^-1^ K_2_HPO_4_, 0.4 g L^-1^ KH_2_PO_4_, 0.1 g U^1^ MgSO_4_·7H_2_O, 0.1 g L^-1^ CaC1_2_·2H_2_O, 4.0 g L^-1^ NaHCO_3_, 4.0 g L^-1^ porcine gastric mucin (cat# M1778, Sigma-Aldrich), 0.25 g L^-1^ sodium cholate and 0.25 g L^-1^ sodium chenodeoxycholate.

In this study, a stool sample was obtained from a healthy volunteer (age 31 years; male). The participant signed informed consent and the sampling protocol was approved by the Ottawa Health Science Network Research Ethics Board at the Ottawa Hospital (#20160585-01H). Stool was collected in pre-reduced sterile PBS buffer, and was immediately homogenized and filtered with sterile gauze in an anaerobic workstation (5% H_2_, 5% CO_2_, and 90% N_2_ at 37 °C). The filtrate was aliquoted in 1.5 ml Eppendorf tubes and stored at −80 °C.

When culturing, the frozen inoculums were taken out and placed in a 37 ° C water bath for 10 minutes, immediately inoculated at a concentration of 2% (w/v) into a 96-well deep well plate containing 1 ml culture medium. Concentration of metformin (metformin hydrochloride) was determined based on the assumption that maximal oral dosage of the drug distributed in 200 g average weight of the colon contents, i.e. 5 μl filter-sterilized water solution of metformin was added into each of the designated wells to achieve a final drug concentration of 7.6 mM^26^. All samples are cultured for optical density and metaproteomics analyses in technical triplicates at each different time point.

### Growth curve fitting with logistic equation

A biobanked human gut microbiome sample were inoculated into the MiPro medium without metformin for culturing, triplicates of wells were collected at 0 (immediately after inoculation), 1.5, 3, 5, 6.5, 8, 10, 12, 13.5, 32, and 48 hr, respectively. Optical density of each sample was measured at 595 nm (OD_595_), as a proxy of microbial biomass. Briefly, 100 μl aliquots were removed from each well, centrifuged at 16,000 ×*g* and the supernatant was used as the medium blank for the OD_595_ measurement. The growth curve is fitted with a logistic growth model by the R package Growthcurver based on the OD_595_ value over 48 hours^36^. The logistic equation is a common standard form in ecology and evolution^37, 38^, whose parameters (the growth rate, the initial population size, and the carrying capacity) provide meaningful population-level information with straight-forward biological interpretation. For comparing the effect of stimulation times on microbiome response to metformin, we determinate metformin was added at the 0 hr of the lag phase, 5 hr of the log phase and 12 hr of the stationary phase, respectively, according to the growth curve. Then, following each metformin treatment, the samples in each group were collected at collected at 0 (immediately after inoculation), 1.5, 3, 5, 6.5, 8, 10, 12, 13.5, 15, 17, 20, 24, 29, and 36 hr for microbial biomass analysis, respectively, as shown in Supplementary Table S2.

### Metaproteomic sample processing and LC-MS/MS analysis

Following each metformin treatment, the samples were collected at 0, 1.5, 5, 6.5, 10, 12, 13.5, 17, 24, 29, and 36 hr for metaproteomic analyses, respectively, as shown in Supplementary Table S2. The collected samples were washed twice by PBS with centrifugation at 14,000 g, 4 °C for 20 min. Then this pellet fraction was then harvested for further metaproteomics analysis. Protein extraction and trypsin digestion were performed according to our previously published procedure^39^. Briefly, the microbial cell pellets were lysed with protein lysis buffer containing 4 % (w/v) sodium dodecyl sulfate (SDS), 8 M urea in 50 mM Tris-HCl buffer (pH 8.0), Roche PhosSTOP™ and Roche cOmplete™ Mini tablets. Lysates were sonicated with a Q800R3 sonicator (QSonica) for 5 min at 4 °C (60 secs on/off, 70% amplitude). Then Removing cell debris through high-speed centrifugation at 16,000 g, 4 °C for 10 min. The resulting supernatant was precipitated using acidified acetone/ethanol buffer at −20 °C overnight.

After centrifugation at 16,000 g for 20 min at 4 °C, the pelleted proteins were washed three times with ice-cold acetone, then dissolved in 6 M urea in mM ammonium bicarbonate (pH = 8). Then, 50 μg of protein in each sample was reduced and alkylated with 10 mM dithiothreitol and 20 mM iodoacetamide, respectively. One microgram of trypsin (Worthington Biochemical Corp., Lakewood, NJ) was then added and shaken for digestion at 37 °C overnight. Protein concentrations were determined by DC (detergent compatible) protein assay.

The tryptic peptides were desalted with a 10-μm C18 column and desalted tryptic peptides corresponding to 0.2 ug of protein were loaded for LC-MS/MS analysis with an Eskigent 425 nanoLC and run on a Q Exactive HF-X mass spectrometer (ThermoFisher Scientific Inc.). Peptides were separated with an analytical column (75 um i.d. × 15 cm) packed with reverse phase beads (1.9 μm; 120-Å pore size; Dr. Maisch GmbH, Ammerbuch, Germany). Peptide separation was performed using a 90 min gradient from 5 to 35% (v/v) acetonitrile at a flow rate of 300 nl/min. The instrument parameters included a full MS scan from 300 to 1800 m/z, followed by data-dependent MS/MS scan of the 16 most intense ions, a dynamic exclusion repeat count of two, and repeat exclusion duration of 30 s. All samples were run on LC-MS/MS in a random order. In addition, nine samples were randomly selected for MS QC runs.

### Metaproteomics data processing

The MS data was processed with MetaPro-IQ bioinformatics workflow^39^ by the MetaLab software (version 1.0)^21^ for peptide/protein identification and quantification. Database construction were based on an iterative database search strategy using gut microbial gene catalogs (from http://meta.genomics.cn/) and a spectral clustering strategy. The identified protein lists were generated with a target-decoy strategy based on a FDR cut-off of 0.01. Label-free quantification (LFQ) intensity of proteins across all samples were obtained with the MaxLFQ algorithm^17^. Taxonomic and functional enrichment analysis was performed using the enrichment module on iMetalab.ca^40^ through inputting the list of altered proteins groups. The database (version 2014) of clusters of orthologous groups (COG) of proteins was used for functional annotation^39^.

### Statistical and multivariate data analysis

In order to compare the difference among the different groups in microbial biomass and protein groups at each time point, the data at the missing time points were imputed using the R package imputeTS^41^, which specializes on univariate time series imputation. The quantified protein groups were filtered with the criteria that the protein should be present in ≥25% of the samples (Q25). And one sample were screened out due to ineligible sample quality (<1/2 average peptide identification). Then LFQ protein group intensities of the filtered file was log2-transformed and normalized through quotient transformation (x/mean) using the MetaboAnalyst (http://www.metaboanalyst.ca/)^42^, as shown in Supplementary Table S7.

To compare the difference in growth curve and metaproteomic profiles between different treatment times, t-test was performed at the time points for 1.5-5 hr (control vs. Lag only), and one-way ANOVA was performed at the time points for 6.5-36 hr. The metaproteomics data was time-course data, involving different treatments (added metformin at different growth phase as a different treatment) and sampling time points. For the time-series analysis, ANOVA-Simultaneous Component Analysis (ASCA) was used to identify major patterns with regard to the two given factors (Treatment and Time) and their interaction (Treatment versus Time). The algorithm first partitions the overall data variance (X) into individual variances induced by each factor (Treatment and Time), as well as by the interactions (Treatment versus Time)^43^. The formula is shown below with (E) indicates the residual Errors: X = Treatment + Time + Treatment versus Time + E. To test the significance of the effects associated with main effects, we performed a model validation, which is based on the Manly’s unrestricted permutation of observation then calculate the permuted variation associated with each factor. In general, protein groups with high leverage exhibit and low squared prediction error (SPE) value in ASCA analysis, which means that significant model contributions are also well modeled protein groups^44^. Cut off values for SPE and leverage were computed as described in the methods section taking α=0.05. The t-test, ANOVA, ASCA analysis and Principal component analysis (PCA) was performed in MetaboAnalyst 4.0^45^. Hierarchical clustering analysis and heatmap plotting was performed with Perseus (version 1.6.10.0)^46^. Multivariate analysis of variance (MANOVA) was performed in MATLAB (The MathWorks Inc.).

## Acknowledgments

This work was supported by the Government of Canada through Genome Canada and the Ontario Genomics Institute (OGI-156), the Natural Sciences and Engineering Research Council of Canada (NSERC, grant no. 210034), and the Ontario Ministry of Economic Development and Innovation (ORF-DIG-14405).

## Author contributions

Z.H., L.L., H.L., and D.F. designed the study. D.F. supervised the study. Z.H, L.L., Z.N., and K. W. performed the experiments. Z.H, L.L., Z.N., X. Z. and K.C. performed data analysis. Z.H., L. L., and D.F. wrote the paper. H.L., X.Z., J.M., K.W., and Z.N. contributed to the editing and revision of the paper. All authors read and approved the final manuscript.

## Disclosure of Potential Conflicts of Interest

No potential conflicts of interest were disclosed.

